# Seeding patient-derived tau induces tauopathy-specific aggregation and lysosomal disruption in human cells

**DOI:** 10.64898/2026.04.20.719763

**Authors:** Tomas Kavanagh, Aysha Strobbe, Kaleah Balcomb, Charleigh Agius, Jingting Gao, Sian Genoud, Evgeny Kanshin, Beatrix Ueberheide, Michael Kassiou, Eryn Werry, Glenda Halliday, Eleanor Drummond

## Abstract

**Background:** Tau aggregation is the defining feature of tauopathies, however, the mechanisms by which distinct tau strains drive disease-specific responses remain unclear. Existing models largely rely on recombinant tau seeding or tau overexpression, which fail to capture the biochemical diversity of pathological tau. The aim of this study was to develop a robust and reproducible human cell-based model of disease-specific tau pathology and to use this model to determine how tau from unique diseases impact tau accumulation and lysosomal dysfunction.

**Methods:** Patient-derived tau aggregates were enriched from post-mortem brain tissue obtained from sporadic Alzheimer’s disease (AD), Pick’s disease (PiD), progressive supranuclear palsy (PSP), and control cases using phosphotungstic acid precipitation. Patient-derived tau preparations were biochemically characterised by immunoblotting and mass spectrometry and normalised for tau content prior to seeding. Patient-derived tau aggregates were seeded into multiple human immortalised cell lines (SH-SY5Y, M03.13, U-87 MG, and U-118 MG cells) and iPSC-derived astrocytes. Tau seeding efficiency, aggregate morphology, and integrity of the autophagy–lysosomal pathway was assessed using quantitative imaging approaches.

**Results:** Patient-derived tau seeds retained disease-specific phosphorylation patterns and isoform composition and led to reproducible, dose-dependent insoluble tau accumulation in all cell lines tested. Despite equivalent tau input and similar background protein composition, PiD-derived tau had the most aggressive pathological signature, showing the highest number of tau aggregates per cell and inducing system wide disruptions in the autophagy lysosomal system including increased SQSTM1 puncta and lysosomal damage markers. Seeding with AD-derived tau led to a high number of tau aggregates per cell and more specifically depleted the lysosomal protease CTSD and uniquely co-seeded Aβ pathology. Seeding with PSP-derived tau resulted in only a moderate number of tau aggregates per cell and uniquely caused increased lysosomal biogenesis.

**Conclusions:** Together, these results demonstrate that intrinsic properties of human tau strains drive disease-specific cellular responses and establish a scalable, physiologically relevant platform for dissecting tau-cell interactions and screening therapeutics across tauopathies.

## Background

Tauopathies are a diverse group of neurodegenerative diseases characterised by the pathological aggregation of tau. There are over 20 clinical diseases driven by tauopathy, which account for a substantial proportion of neurodegenerative diseases featuring cognitive and motor decline [1–4]. Tauopathies can be categorised based on the tau isoform that preferentially aggregates in disease: 4R in progressive supranuclear palsy (PSP), 3R in Pick’s disease (PiD) and a mixture of 3R+4R in Alzheimer’s disease (AD) [5, 6]. Whilst tau aggregation is a pathological hallmark, tau itself significantly differs in biochemical, structural and cellular features in each of the diseases [5, 7–12]. Distinct isoform mixtures (3R, 4R and 3R+4R), post-translational modifications and filament conformations give rise to disease-specific strains of tau that likely contribute to clinical and mechanistic heterogeneity [13–16]. How these divergent molecular properties of tau alter tau toxicity, aggregation, and cell-type vulnerabilities remains poorly understood.

There is mounting evidence that suggests tau pathology is able to spread between cells in a prion-like manner [17–20]. The ‘seeding’ capacity of tau has been utilised to generate cell and animal models of tau pathology. However, a major limitation of these models is their reliance on synthetic tau fibrils or genetically engineered tau overexpression to establish pathology, which does not recapitulate the disease-specific tau aggregates observed in patients [13, 19–30]. Mouse models of tauopathy frequently require multiple genetic manipulations, are time-consuming to generate, and develop only some aspects of patient pathology [22, 31]. Recent proteomics and structural biology studies now suggest a diverse and complex pool of tau species [13, 16, 32–36]. Accurate modelling of this complexity is not feasible with overexpression studies alone. Patient-derived seeding models have shown promise in replicating tau seeding whilst maintaining the biochemical properties of the unique pools of tau in different tauopathies [37]. However, to date, most studies have focused on the dynamics of tau seeding in bio-reporter cell lines or used cell and animal models already overexpressing wild-type or mutant tau to help drive aggregation [29, 38–41]. Attempts to seed patient-derived tau pathology into cells and animals have been limited by multiple factors: inconsistent biochemical characterisation, low reproducibility, variable seeding and pathology induction, and a lack of appropriate controls or understanding of co-purified proteins [22, 23, 30, 37, 42]. These are critical shortcomings that need to be addressed for models of human tau seeding to become a reliable and reproducible model for the field.

The aim of this study was to develop human cell models that reproducibly express the complex pools of tau present in the brain in sporadic cases of three human tauopathies: AD, PSP and PiD. To do this, we seeded multiple human neuronal and glial immortalized cell lines (SH-SY5Y, M03.13, U-87 MG, and U-118 MG cells) and iPSC-derived astrocytes without tau overexpression with insoluble proteins extracted directly from human AD, PSP and PiD brain tissue. We show that these patient-derived tau aggregates retain their biochemical properties and elicit disease-specific responses in the autophagy-lysosomal system. By including tau derived from control cases and normalising for tau input between cases we provide a means for directly assessing the unique effects of tau in each tauopathy. This model provides a physiologically relevant framework for studying tau interactions, the mechanisms by which unique tau strains drive disease processes and is flexible and scalable enough to allow for medium-throughput drug screening.

## Materials and Methods

### Human brain tissue

All procedures were performed under protocols approved by the Sydney University Human Ethics Committee. In all cases, written informed consent for research was obtained from the patient or legal guardian, and the material used had appropriate ethical approval for use in this project. All patients’ data and samples were coded to protect patients’ identities. All tissue used was acquired from the Sydney Brain Bank, Australia. For patient-derived tau preparations, mass spectrometry, western blot and dot blot, we used fresh frozen tissue taken from the superior frontal cortex at the level of the head of the caudate nucleus from *n* = 3 patients with AD, *n* = 3 patients with PiD, *n* = 3 patients with PSP and *n* = 1 age-matched control. Primary inclusion criteria included neuropathological diagnosis of AD, PSP, PiD or no neurodegenerative disease based on the presence of tauopathy-specific pathology as determined by histological assessment performed by neuropathologists at the Sydney Brain Bank. Cases with no or minimal co-pathology were prioritised to the best of our ability (e.g., no or minimal AD neuropathologic change in tauopathies, TDP-43 pathology, Lewy bodies, aging-related tau astrogliopathy (ARTAG), primary age-related tauopathy (PART), hippocampal sclerosis, and small vessel disease). Cases were selected based on neuropathology alone: clinical diagnosis was not considered in case selection but is provided in Table 1.

**Table 1:** Case demographics for all tissue used in this study. P.M.D = post-mortem delay; bvFTD = behavioural variant frontotemporal dementia; FTD = frontotemporal dementia.

| *Case* | *Age* | *Sex* | *P.M.D. (hrs)* | *ABC* | *Clinical*<br>*Diagnosis* |
| --- | --- | --- | --- | --- | --- |
| *CTRL* | 79 | M | 8 | A0B1C0 | Asymptomatic |
| *AD 1* | 75 | F | 14 | A3B3C3 | AD |
| *AD 2* | 70 | M | 23 | A3B3C3 | AD |
| *AD 3* | 73 | M | 26 | A3B3C3 | AD |
| *PiD 1* | 74 | M | 22 | A0B0C0 | bvFTD |
| *PiD 2* | 73 | M | 11 | A0B0C0 | bvFTD |
| *PiD 3* | 66 | F | 22 | A0B1C0 | bvFTD |
| *PSP 1* | 70 | F | 16 | A0B1C0 | FTD |
| *PSP 2* | 67 | M | 15 | A0B2C0 | FTD |
| *PSP 3* | 70 | M | 18 | A0B1C0 | FTD |

### Tissue homogenization

Tissue homogenization was performed as per previously published methods [19, 24, 38, 43, 44]. Briefly, 250 mg of grey matter was dissected from frozen tissue blocks of each sample. The tissue was then pulverised on dry ice with a hammer. Pulverised tissue was homogenised in low salt homogenization buffer (50 mM HEPES pH 7.0, 250 mM sucrose, 1 mM ethylenediaminetetraacetic acid [EDTA]), protease inhibitor cocktail (cOmplete mini tablets, EDTA-free Millipore Sigma), and phosphatase inhibitor cocktail (PhosSTOP EASYpack, Roche) with a Dounce homogeniser. Protein concentration was then determined using a Micro bicinchoninic acid (BCA) assay kit (Thermo Micro BCA Assay Cat. No.: 23235). Homogenised samples were flash-frozen in a dry ice:ethanol slurry and stored at −80°C.

### Phosphotungstic acid precipitation and enrichment of tau

Phosphotungstic acid (PTA) precipitation was performed as described [32, 45, 46]. Briefly, 20% (w/v) brain homogenate was mixed with sarkosyl detergent and benzonase to a final concentration of 2% (v/v) and 0.5% (v/v), respectively. Samples were incubated for 2 hr with constant shaking (1,200 rpm) at 37°C in a ThermoMixer (Eppendorf). Next, 10% w/v Sodium phosphotungstic acid solution (pH 7.0) was added to a final concentration of 2% (v/v) and incubated overnight at 37°C with constant shaking (1,200 rpm). The sample was centrifuged at 16,000*g* for 30 min at room temperature, and the supernatant was removed. The resulting pellet was resuspended in 2% (v/v) sarkosyl and 2% (v/v) PTA in double-distilled H_2_O, pH 7.0. The sample was again incubated for at least 1 hr with constant shaking (1,200 rpm) at 37°C before a second centrifugation as above. The supernatant was again removed, and the pellet was resuspended in PBS using 10% of the initial starting volume.

### Biochemical analysis of patient-derived tau seeds

Dot blots were performed to determine the relative enrichment of tau in patient-derived tau preps. Samples were mixed 1:1 with 16 M urea and 2% SDS for 10 minutes using 2 µL of sample per dot. BSA (bovine serum albumin) was dotted as a negative control. The total volume was then dotted onto a nitrocellulose membrane 2 µL at a time and allowed to dry for 1 min before blocking with 5% skim milk or 5% BSA in Tris-buffered saline with Tween-20 (TBS-T) for 1 hr. Blots were incubated for 2 hr at room temperature with primary antibodies for total tau (Invitrogen, MA5-12808, 1:1000), pTau217 tau (Invitrogen, 44-744, 1:1000), AT8 tau (Invitrogen, MN1020, 1:1000), pTau396 tau (Abcam, #ab109390, 1:1000 and Cell Signalling Technology, 9632 [PHF13], 1:1000), Amyloid-β (4G8, Biolegend, 800701, 1:1000), α-synuclein (BD, 610787, 1:1000), pTDP^432A^ (Cosmo Bio, CAC-TIP-PTD-P07, 1:1000), pTau422 (Abcam, #ab79415, 1:1000), T22 (Merck, ABN454-I, 1:1000), or β-actin (AC-15, Abcam, #ab6276). Dot blots were then incubated for 2 h at room temperature with secondary antibody (anti-rabbit, anti-mouse or anti-goat horseradish peroxidase (HRP) conjugated antibody, 1:10,000). Western blots were then developed with electrochemiluminescence (ECL) Western blotting substrate (Merck #WBULS0500) and imaged on a Thermo Scientific CL1500 gel imaging system. To determine the concentration of tau in each PTA preparation, a total tau ELISA was used (Invitrogen, KHB0041). Samples were diluted 1:1 with 6 M urea for 30 minutes before dilution with sample buffer to 1:100. ELISAs were performed as per the manufacturer’s protocols and measured on a BMG ClarioStar plate reader. A 4PL curve was fit to estimate available tau abundance. Samples were then normalised to 10 pg/µL for all tissue culture experiments. 10 µL of the sample was additionally aliquoted for mass-spectrometry analysis.

### Protein solubilization, digestion and LC-MS/MS

Each sample was solubilized in the lysis buffer (5% SDS, 10 mM TCEP 20 mM CAA, and 100 mM TRIS (pH=8). Then all samples were incubated for 30 min at 90 °C (2,000 rpm) and centrifuged at 16,500 × g for 5 min at RT. Supernatants were transferred into 96 well plate for subsequent processing. Protein lysates were supplemented with magnetic SP3 beads, and proteins were precipitated with a 2× fold dilution with EtOH. Proteins captured on magnetic beads were washed 3 times in 85% EtOH before resuspension in 100 ul of 50 mM TRIS (pH=8) containing trypsin. Proteins were digested o/n at 37 °C. Peptides were transferred into a clean 96-well plate, and magnetic beads were washed in 100 ul of 0.1% FA. Peptides were loaded on Evosep One C18 tips for subsequent analysis by LC-MS/MS. MS analysis was performed on the LTQ Orbitrap Eclipse instrument in DIA mode, and RAW MS files were analysed in Spectronaut. The mass spectrometric raw files are accessible at the MassIVE Repository under accession MassIVE MSVxxx. Raw label-free quantitation and IBAQ scores are provided in Supplementary Table 1.

### Tissue culture

SH-SY5Y, M03.13, U-87 MG, and U-118 MG cells were maintained as per ATCC guidelines. All cell lines were submitted for centralised mycoplasma contamination testing quarterly during use, with the Lonza Mycoplasma kit. SH-SY5Y cells were maintained in DMEM/F12 media with glutamax and 10% heat-inactivated fetal bovine serum (FBS). MO3.13 and U-118 MG cells were maintained in DMEM media with glutamax and 10% FBS. U-87 MG cells were maintained in MEM media with glutamax and 10% FBS. All cell lines were maintained at 37°C and 5% CO_2_ in a humidified incubator (Hireseus). Cells were seeded at 5.5 × 10^3^ per well in 96-well plates (cytotoxicity assays), 3.5 × 10^4^ cells per well for 24-well plates (imaging) and 5.5 × 10^4^ cells/well for 12 well plates (protein). Seeded cells were allowed to adhere and recover overnight before treatment with patient-derived tau preparations.

### iPSC derived Astrocytes

iPSC-derived astrocytes (iAstrocytes) were derived in a previous study from healthy control iPSCs sourced from Cedars Sinai iPSC Core Repository (Los Angeles, USA) [47]. This study used Ctrl-06, derived from fibroblasts of an 82-year-old, female healthy control donor (APOE ε3/ε3). Cells were reprogrammed from fibroblasts via episomal plasmids. Their identity as astrocytes were verified using immunofluorescence, functional studies and transcriptomics in comparison to normal human astrocytes [47]. Approval for the use of iPSCs was gained from the University of Sydney Institutional Biosafety Committee and Human Research Ethics Committee.

Upon reaching maturity, iAstrocytes were maintained at 37°C and 5% CO_2_ in a humidified incubator (Thermo Fisher Scientific) and grown until 90% confluent. They were then passaged using Accutase (Sigma) and plated on Matrigel (0.08 mg/well; Corning)-coated glass coverslips in 24 well plates at 9 x 10^3^ cells/well in astrocyte medium (astrocyte basal medium, 2% FBS, astrocyte growth supplement and 10 U/mL penicillin/streptomycin solution; all from ScienCell). After 24 hrs, the astrocytes were treated with tau aggregates, or lipofectamine only.

### Patient-derived tau seeding

Tau seeding experiments were performed by first vigorously vortexing patient-derived tau preparations immediately before use. Patient-derived tau seeds were then mixed with OptiMEM media (Gibco) and lipofectamine-3000 (Thermo) at volumes appropriate to the plate format being used. For 24-well plates, 15, 40, 65 and 105 pg of tau were seeded from normalised aliquots (10 pg/µL tau). This was mixed with 0.75 µL of lipofectamine-3000 in OptiMEM media to a final volume of 50 µL/well. Preparations were incubated at room temperature for 1 hr then seeded dropwise onto cells. For smaller and larger well reactions, quantities were scaled to surface area. For autophagy-lysosomal pathway experiments a control well was treated with 100 µM chloroquine for 1 hr prior to fixation as a positive control for lysosomal disruption.

### Cytotoxicity assays

SH-SY5Y cells were seeded at 5.5 × 10^3^ cells/well. A media change was performed 12 hrs after seeding which included multiplexed RealTime-Glo™ MT Cell Viability Assay (Promega, #G9712) and CellTox™ Green Cytotoxicity Assay reagents (Promega, #G8743). Cells were then treated with PTA tau preparations as above and incubated for 1 hr before the first read. Reads were then performed every 24 hrs for 72 hrs on an Ensight multi-mode plate reader (Perkin-Elmer). Cell toxicity was normalised to viability and plotted for the time course.

### Fluorescent staining

For imaging studies, cells were seeded on 13 mm Φ coverslips at 3.5 x 10^4^ cells per well and grown for four days post-treatment. Cells were fixed for 15 minutes with 4% PFA, followed by 3 washes with PBS. Cells were permeabilised and blocked in 10% BSA + 0.1% saponin for 1 hr at room temperature. Cells were then incubated for 2 hrs at room temperature in primary antibodies diluted in 2% BSA + 0.02% Saponin. Total tau (Tau-5 Invitrogen, MA12-50808, 1:1000), pTau199/202 (Invitrogen, #44-768G, 1:1000), Aβ (Cell Signalling Technology, #8243, 1:1000), LAMP1 (Abcam, ab24170, 1:500), LGALS3 (Invitrogen, #14-5301-82, 1:500), CTSD (R&D Systems, #AF1014, 1:500), p62/SQSTM1 (Abcam, ab56416, 1:500), GFAP (Novus Biologicals, NOVNBP105198, 1:500), and β-III-tubulin (Merck, AB9354, 1:500). Samples were washed by dipping 10 times in 4 containers of PBS. AlexaFluor488-(Jackson ImmunoResearch, #115-545-164, 1:500), CF568-(Sigma, SAB4600036, 1:500), AlexaFluor647-(Invitrogen, A32787, 1:500), conjugated secondary antibodies or CellMask Deep Red dye (Invitrogen, C10046, 1:1000), were incubated for 1 hr at room temperature with Hoechst 33342 (Sigma, B2261, 1:2000). Coverslips were washed as above and then mounted onto glass slides with Prolong Glass Antifade Mountant (Invitrogen, P36984) and allowed to set for 72 hrs before imaging.

### Imaging

Widefield images were taken with a Leica Thunder imaging system using a 63x (NA 1.4) objective, tiling five 3×3 tiles with a 10% smoothed overlap without deconvolution on each coverslip. Confocal images were captured at 60x (NA 1.4) on a Nikon C2 Confocal microscope or at 63x (NA 1.4) on a Leica Stellaris 8. All imaging was performed using Leica immersion oil type F. Images were exported as “.tif” images for downstream analysis. Images were processed in ImageJ (v2.14.0). Widefield images were used to quantify tau seeding rates across each treatment. Differences of Gaussians were used to remove background and identify puncta maxima, which were then counted. Simple thresholds and watershedding of Hoechst signal were used to determine cell counts in widefield images. StarDist was used for confocal images. Tau puncta counts were then normalised to cell number. For analysis of tau aggregate size measures, simple thresholds were used to maintain aggregate shapes. Cellpose-SAM was used to create masks of cell bodies to determine the percentage of cells transfected with tau. Quantification of LGALS3 and SQSTM1 puncta was achieved with simple thresholds. CTSD puncta were quantified using differences of Gaussians. Simple thresholds, watershedding and MorphoLibJ’s morphological segmentation plugin were used to quantify LAMP1 lysosome count, size and shape. Image panels were constructed in Adobe Photoshop (v26.4.1), and uniform thresholds were applied to each channel of all images in an experiment. Final figures were made in Adobe Illustrator (v28.1).

### Data processing and statistics

Proteomics data was filtered to remove sparse and non-unique proteins. Filtered datasets were then normalised with the MAD-score as in previous publications [48]. Missing values were imputed via knn method (k = 3). Differential enrichment of proteins was assessed with linear models (∼ 0 + disease) to contrast proteins enriched in each disease and filtered to FDR < 0.05 and |log_2_ FC| > 0.58. Volcano plots were made with the EnhancedVolcano package (v). For imaging data, normality was assessed using Normal Q-Q plots of residuals and distribution histograms for each dataset in combination with Shapiro-Wilk normality tests. Normally distributed experiments were assessed with one- or two-way ANOVA followed by Tukey’s post-hoc test. Mixed-effect linear models were used to assess the variation contributions from biological groups compared to residuals in replication studies with meanTauCounts ∼ disease + (1 | replicate). The VarCorr and fixef functions were used to extract variable correlations (*vcov*), covariate SD (σ), and fixed effects (β). For datasets that were not normally distributed, a Kruskal-Wallis test was performed, followed by Dunn’s post-hoc test. Multiple comparisons were corrected with the Benjamini-Hochberg method. All graphs display the mean ± standard deviation unless otherwise stated. Boxplots represent the median, Q1 and Q3, with whiskers representing 1.5 × IQR. All statistics and graphing were performed in RStudio with R v4.5.2 with the following packages: tidyverse (2.0.0), ggpubr (v0.6.3), ggrepel (v0.9.7), plotrix (v3.8-14), corrplot (v0.95), EnhancedVolcano (v1.28.2), reshape2 (v1.4.5), car (3.1-5), limma (v3.66.0), impute (v1.84.0), lme4 (v1.1-38), lmerTest (v3.2-0), rstatix (0.7.3).

## Results

### Patient-derived tau aggregates include a complex mix of proteins

We enriched tau pathology from *n* = 3 AD, PiD and PSP cases and *n* = 1 control case (Table 1) using sodium phosphotungstate (PTA) precipitation [38, 39] (Figure 1). To determine the relative abundance of disease-associated pathology, we performed dot blots on total homogenate, PTA-enriched fractions, washes 1 (S1) and 2 (S2) and negative control BSA (Figure 2.A). We observed strong enrichment of total tau in the pellet of all cases compared to the control, with PSP showing comparatively lower tau levels than AD and PiD. We observed strong enrichment of insoluble AT8-immunoreactive phosphorylated tau in AD and PiD compared to control, but not in PSP. We did, however, detect pTau217, pTau396, and pTau422 in the PTA insoluble fractions of PSP, albeit at lower levels than observed in AD- and PiD-derived insoluble fractions (Supplementary Figure 1). Oligomeric tau, as assessed by T-22-immunoreactivity, was present in all PTA insoluble pellets, but was only enriched in AD and PiD cases in comparison to the control case (Supplementary Figure 1). We observed enrichment of insoluble Aβ only in AD cases (Figure 2.A). Importantly, we observed no enrichment of other disease-associated pathological proteins, including pTDP43 or α-synuclein, nor of the housekeeping control β-actin (Figure 2.A).

**Figure 1.**
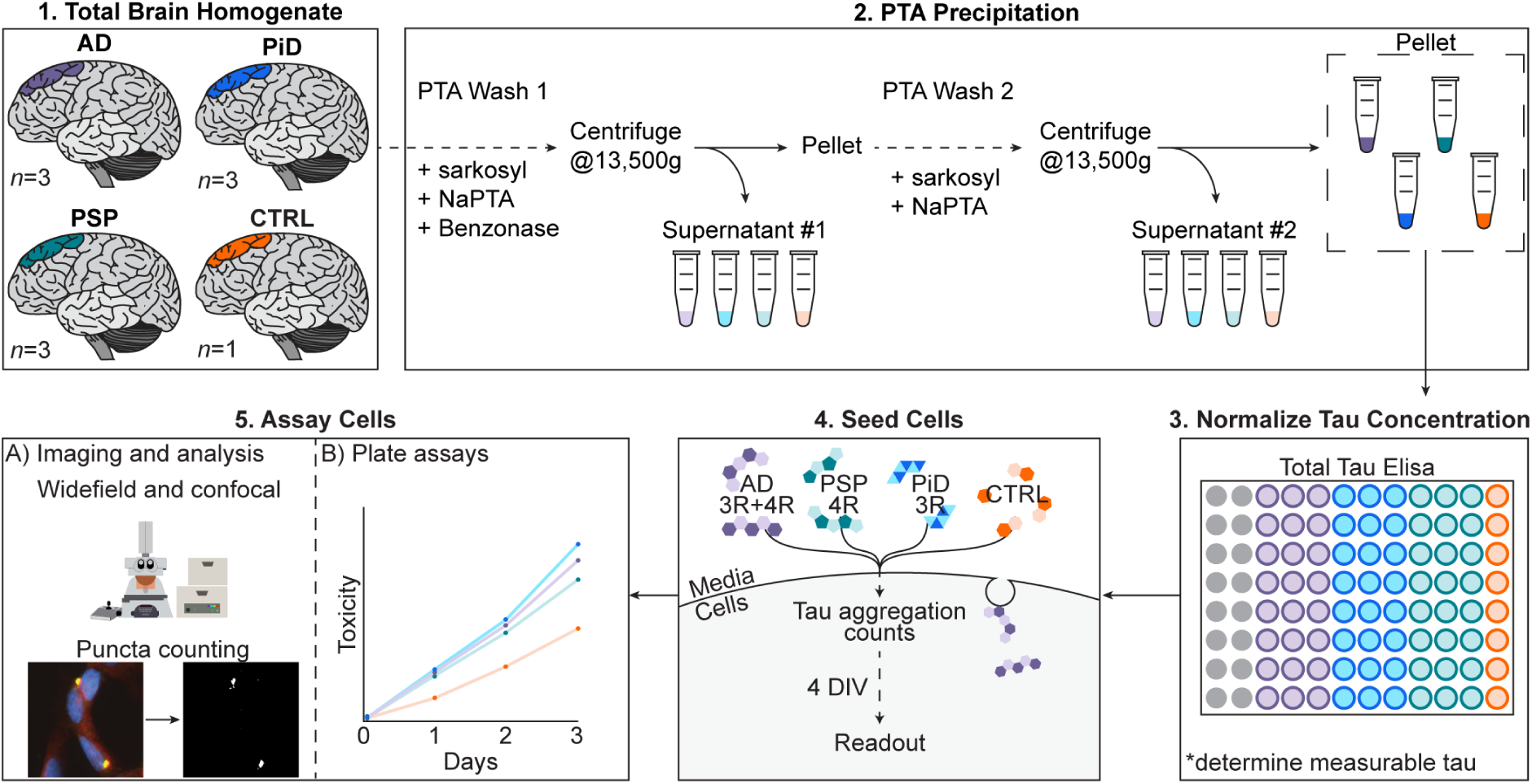
Overview of the experimental workflow

**Figure 2.**
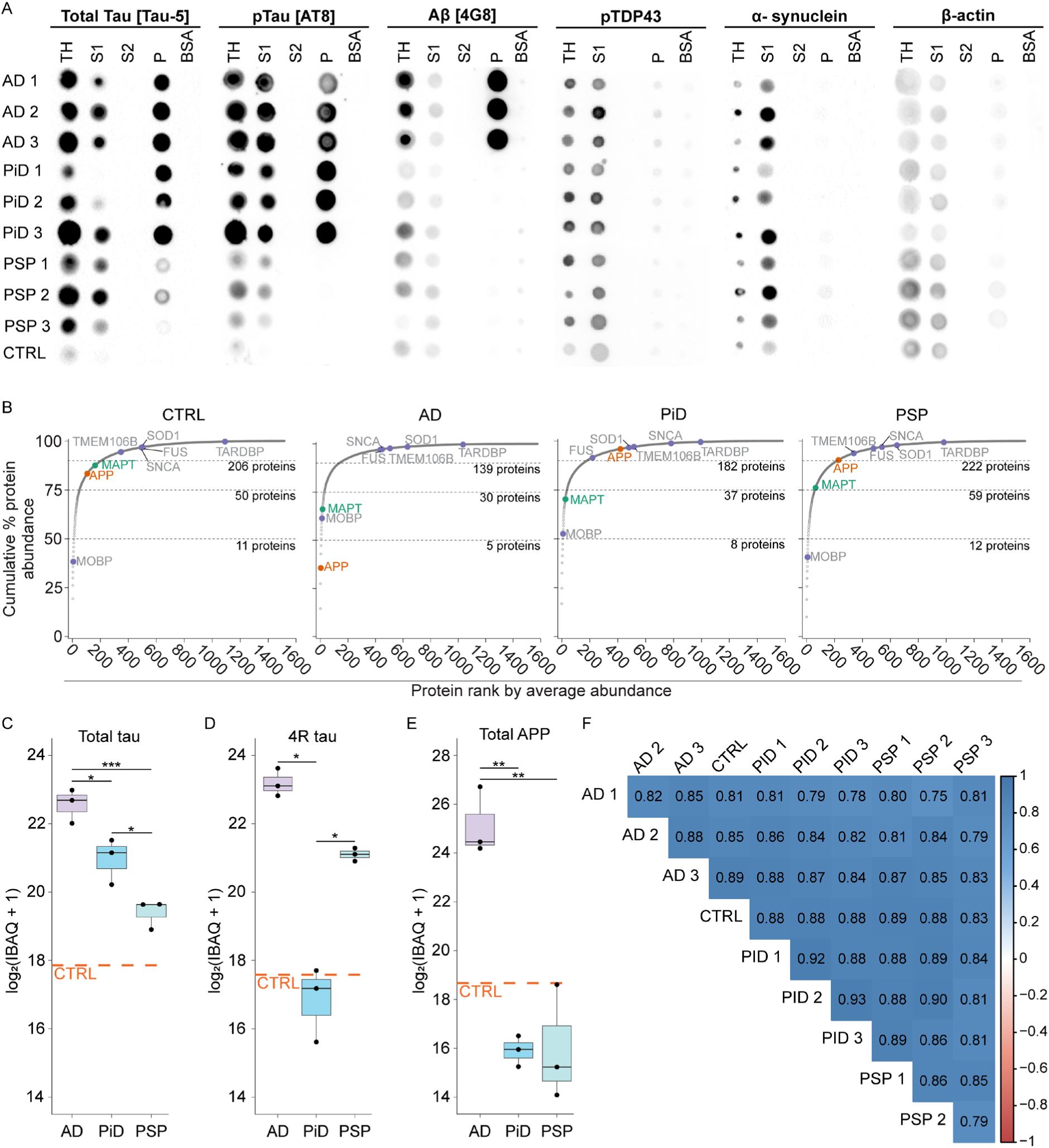
Biochemical analysis of patient-derived tau seeds. A) Dotblots of fractions derived by the PTA enrichment steps probed for total tau [Tau-5], phosphorylated tau [AT8], Aβ [4G8], α-synuclein, phosphorylated TDP-43 and β-actin. B) Cumulative protein abundance contributed to by each protein from all proteins detected. Dashed lines represent 50%, 75% and 90% of total protein abundance cut-offs. C) Total tau abundance measures by MS. Biological *n* = 3 per disease, excluding controls. One-way ANOVA with Tukey’s post-hoc test. Adjusted p-values: PiD-AD = 0.026, PSP-AD = 0.0009, PSP-PiD = 0.028. D) 4R tau specific peptide “LDLSNVQSK” abundance measures by MS. Biological *n* = 3 per disease, excluding controls. One-way ANOVA with Tukey’s post-hoc test. Adjusted p-values: PiD-AD = 0.00006, PSP-AD = 0.022, PSP-PiD = 0.00062. E) APP abundance measures by MS. Biological *n* = 3 per disease, excluding controls. One-way ANOVA with Tukey’s post-hoc test. Adjusted p-values: PiD-AD < 0.001, PSP-AD < 0.001, PSP-PiD = 0.99. F) Correlogram of Pearson’s correlations (r) between all pairwise samples. TH: total homogenate, S1: supernatant 1, S2: supernatant 2, P: pellet, BSA: bovine serum albumin, CTRL: Control case. Boxplots represent the median, Q1 and Q3, with whiskers representing 1.5 × IQR.

To more deeply profile the PTA-insoluble fraction enrichment from all cases we performed mass spectrometry. We detected 1,592 proteins in total, with the vast majority at very low intensities (88% of proteins represent <10% of total protein abundance). This analysis revealed tau peptides accounted for 0.06-1.8% of total spectra (Figure 2.B, C, Supplementary Table 1, 2). AD had the highest degree of tau enrichment, with an average abundance rank of 16 and an average total protein percentage of 1.13 ± 0.58%. PiD had the second strongest enrichment of tau, with an abundance rank of 26 and an average total protein contribution of 0.46 ± 0.19%. Tau in PSP had an abundance rank of 65 and an average total protein contribution of 0.20 ± 0.07%. In control, tau was ranked 162, representing 0.06% of total protein. Tau isoforms biases were maintained in all preparations with PiD having considerably less 4R tau enrichment (Figure 2.D). As with the dot blots, APP (Aβ) was only enriched in AD (abundance rank 3, 7.89 ± 7.2% of total protein) and was in the bottom 10% of proteins for all other cases. APP abundance was low in PiD (rank 419, 0.01 ± 0.01% of total protein) and PSP (rank 229, 0.04 ± 0.05% of total protein). In the control sample, APP was ranked 106, representing 0.1% of total protein.

Proteins associated with other disease pathologies were predominantly in the bottom 10% of proteins (e.g. α-synuclein, FUS, TDP-43, TMEM106B, SOD1, Figure 2.B) and contribute little to overall protein composition. Between 30 and 59 proteins accounted for 75% of the total pool of insoluble proteins. Critically, the control case closely mimicked the enrichment profiles of background proteins seen in disease cases. This included the enrichment of the most abundant off-target proteins – MBP, MOBP, COL1A2, H1.1, H1.2, SAP. In fact, the overall protein composition of each sample was highly correlated (Pearson’s r range 0.75 – 0.93, Figure 2.F). This highlights the importance of using a control brain sample to account for the contaminant protein effects seeded into cells in these experiments.

Finally, we assessed that few proteins contributed to the bulk of variation between each sample. We used standard deviation to find proteins with high variation across all samples and differential expression analysis to determine proteins differing between groups. Most variation between cases was driven by 44 proteins (Supplementary Table 1). The most variable proteins included PSMC5, FMN2, SLIT2, ACOT2 and APP (Supplementary Table 1, Figure 2.E). Many proteins associated with Aβ plaques contributed significant variability across groups, including APOE, MDK and SMOC1 (SD = 3.3, SD = 2.1 and SD = 2.0, respectively) [49, 50]. Tau variation across all samples was relatively low (SD = 1.57), suggesting relatively stable enrichment of tau in disease samples. This is supported by differential expression analysis that showed minimal differential expression between samples. Only contrasts with AD showed significant results (10 proteins in AD vs PiD, 16 proteins in AD vs PSP). There were no significant differences between PiD- and PSP-derived tau preparations. These analyses identified that most of the variation was driven by higher enrichment of proteins in AD samples (Supplementary Table 3, 4, Supplementary Figure 2). Some examples of tauopathy-specific contributions to overall variation include CALML5 which shows higher enrichment in PiD samples. This analysis suggests a relatively small proportion of all proteins enriched by the PTA protocol are driving most of the variation.

### Patient-derived tau aggregates seed SH-SY5Y cells without tau overexpression

To determine if patient-derived tau aggregates could seed pathology in cells without tau overexpression, we seeded increasing doses of tau seeds into wild-type SH-SY5Y cells. This cell line was chosen because it is commonly used as an immortalised neuronal-like cell line in neurodegenerative disease studies, is amenable to manipulation with simple genetic tools (plasmids, siRNAs), and can be used in rapid screens. We seeded 15, 40, 65 and 105 pg of tau from each case (n=3 AD, n=3 PiD, n=3 PSP and n=1 control) into 24-well plates containing SH-SY5Y cells (Figure 3.A). In all cases, we saw successful seeding of tau pathology at the 4-day timepoint imaged (Figure 3.A). Furthermore, tau seeds had minimal impact on cell toxicity/viability when treated with the same dose-response curve (Figure 3.B). We saw no difference between treatment groups up to 72 hrs post-treatment, suggesting that WT SH-SY5Y cells may experience little seeding-induced disease-specific toxicity (Figure 3.B). We observed robust seeding of tau aggregates in all disease-derived cases.

**Figure 3.**
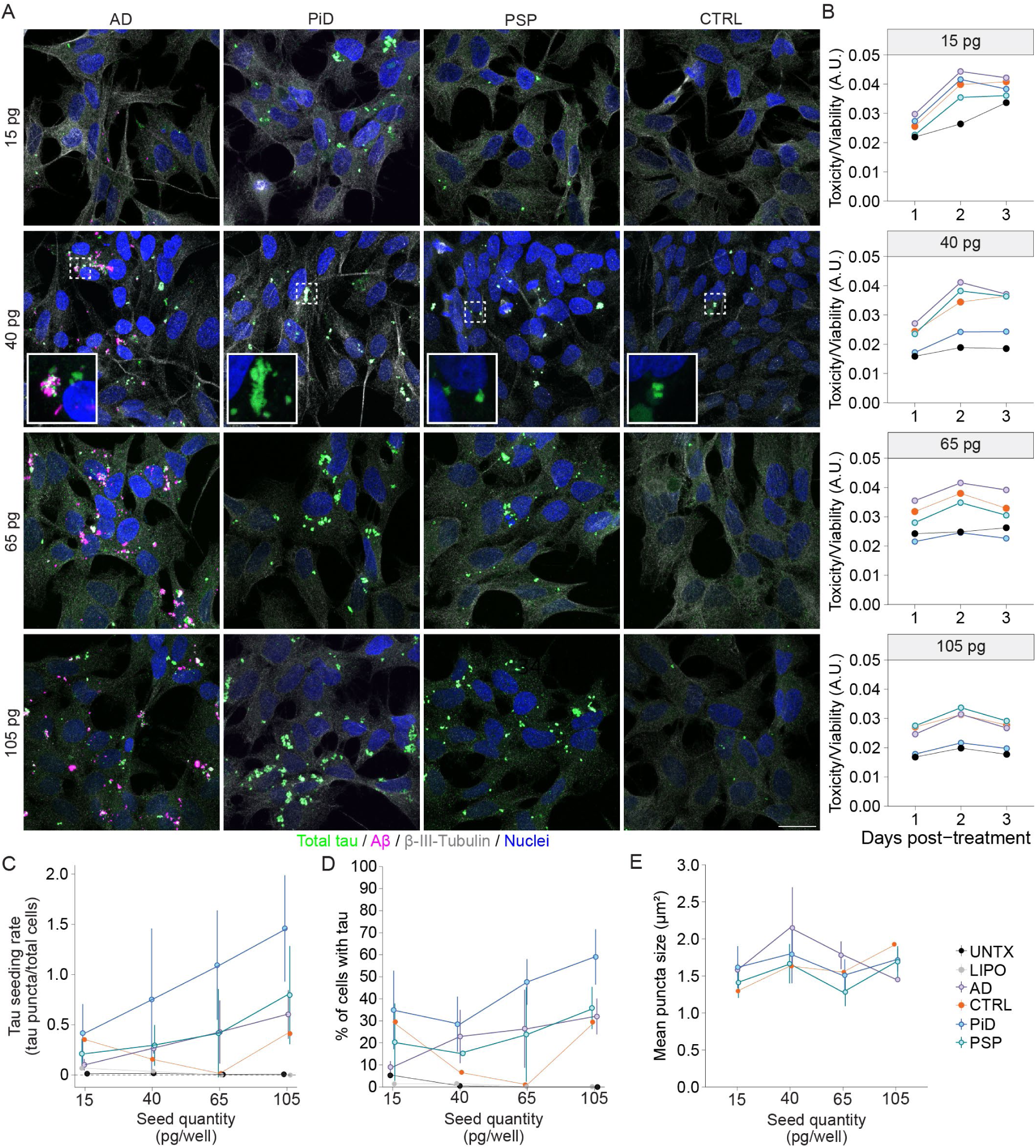
Patient-derived tau seeds follow a linear dose-response. A) Representative confocal images of SH-SY5Y cells seeded with 15, 40, 65 and 105 pg of patient-derived tau. B) Cyto-toxicity responses of SH-SY5Y cells to treatment with patient-derived tau seeds at each concentration. C) Quantification of tau seed number dose-response to patient-derived tau seeds. Two-way ANOVA followed by Tukey’s post-hoc test. Disease effect F = 7.87 and adjusted p-value = 0.0023, treatment quantity F = 5.49, adjusted p-value = 0.005. D) Percentage cells transfected with tau at each seeded concentration. Two-way ANOVA followed by Tukey’s post-hoc test. Disease effect F = 7.53 and adjusted p-value = 0.0029, treatment quantity F = 4.40, adjusted p-value = 0.013. E) Tau aggregate size changed with different dosing levels of tau. Two-way ANOVA followed by Tukey’s post-hoc test. Disease effect F = 2.36 and adjusted p-value = 0.12, treatment quantity F = 3.36, adjusted p-value = 0.036. Scale bar = 20 μm. A.U. = arbitrary units. *N = 3* biological replicates for AD, PiD and PSP, *n = 1* CTRL. LIPO = lipofectamine-only treatment, UNTX = untreated. Line plots are mean ± standard deviation.

We determined the dose-specific seeding efficiency by quantifying tau aggregates normalised to total cells at 4 days post-treatment. PiD-derived tau resulted in maximal seeding, generating between 0.41 ± 0.29 – 1.46 ± 0.53 aggregates per cell at 15-105 pg treatments (Figure 3.C). AD gave the second-highest seeding rate with 0.1 ± 0.06 to 0.6 ± 0.24 tau aggregates/total cell count. Tau puncta/total cell count for control-derived tau-treated cells ranged from 0.01 – 0.41 (*n =* 1, Figure 3.C). PSP-derived tau led to 0.21 ± 0.15 to 0.8 ± 0.49 tau puncta/total cell count (Figure 3.C). In cells seeded with AD-derived tau, we also observed Aβ aggregates (Figure 3.A), which showed considerable variability between cases as treatments were normalised to tau quantity (Supplementary Figure 3.A). We observed negligible tau aggregation in cells treated with lipofectamine only or no treatment (Figure 3.C, Supplementary Figure 3.B). Tau aggregates tended to cluster near nuclei and were typically homogenous in texture and intensity (Figure 3.A). Increasing tau concentration increased the seeding rate (Figure 3.C, treatment p-value = 0.005, F-value = 5.5), while exerting a modest effect on overall cell transfection efficiency (Figure 3.D, treatment p-value = 0.02, F-value 3.9). This indicates that increasing seeding quantity primarily increases the number of tau aggregates taken up by an individual cell rather than increasing the total number of cells with tau aggregates. However, the disease origin of patient-derived tau aggregates had a strong effect on the percentage of cells seeded with tau pathology (disease p-value = 0.0002, F-value = 10), with PiD-derived tau seeding from 28.5 - 59% of cells (Figure 3.D). AD and PSP had similar percentages of cells with tau aggregates (AD: 9 - 32%, PSP: 15 - 36%). Tau aggregate size did not change with seeding concentration (Figure 3.E) or disease. Together, these results suggested that seeding with 40 pg of tau was optimal for subsequent experiments in SH-SY5Y cells, as higher concentrations did not sufficiently enhance seeding to justify the increased use of human brain tissue.

### Patient-derived tau consistently seeds aggregates in SH-SY5Y cells

To determine the reproducibility of tau seeding in SH-SY5Y cells, we repeated these experiments three times for 15 and 40 pg quantities of patient-derived tau seeds. We applied mixed-effects models to evaluate the relationship between replicate-level variance (*vcov*_rep_), residuals (*vcov*_res_) and the signal (β) of each PTA-derived tau seeding rate. At 15 pg seeding rates suffered from low effect sizes and high variability (*vcov*_rep_ = 0.031, *vcov*_res_ = 0.082, β = - 0.254 – 0.057), with residual variances dominating actual effects in linear models (Supplementary Table 5). As such, variation in one replicate could easily skew the overall experiment (Figure 4.A, B). Despite the variability in tau seeding rate after seeding 15 pg, the size of tau aggregates remained largely stable between replicates (Figure 4.C). However, transfection efficiency was very variable at 15 pg treatment (Figure 4.D).

**Figure 4.**
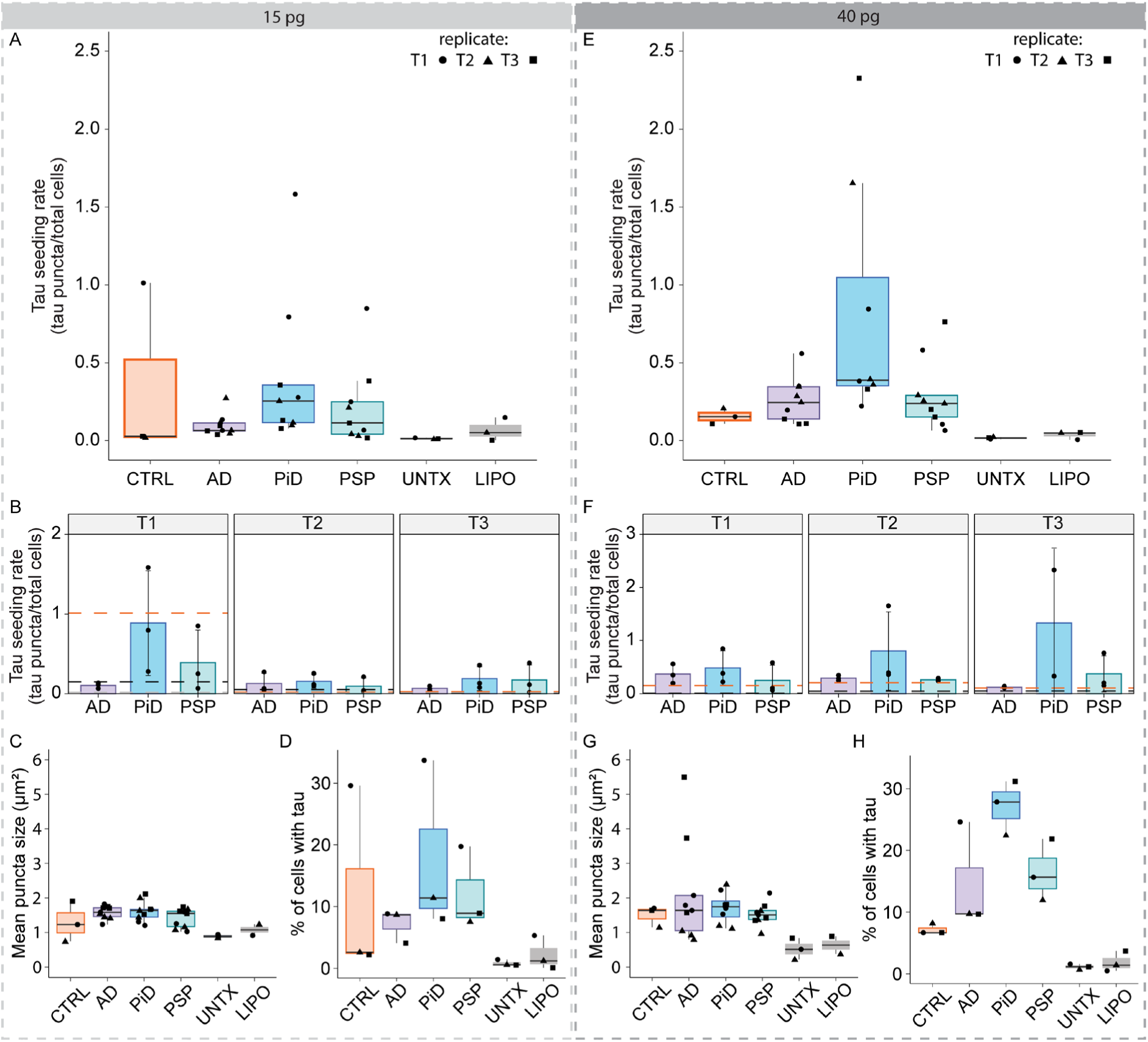
Tau seeding is replicable at 40 pg. 15 pg treatments of tau patient-derived tau seeds across three technical replicates A-D. A) Boxplot representing tau puncta counts normalised to total cell count across three technical replicates of each disease-derived tau preparation treated with 15 pg. B) Tau puncta count of each technical replicate with 15 pg treatment. C) Average tau puncta size across technical replicates with 15 pg treatment. D) Percentage of cells transfected by 15 pg of tau seeds across three technical replicates. 40 pg treatments of tau patient-derived tau seeds across three technical replicates E-H. E) Boxplot representing tau puncta counts normalised to total cell count across three technical replicates of each disease-derived tau preparation treated with 40 pg. F) Tau puncta count of each technical replicate with 40 pg treatment. G) Average tau puncta size across technical replicates with 40 pg treatment. H) Percentage of cells transfected by 40 pg of tau seeds across three technical replicates. T1/2/3 = technical replicate 1/2/3.

For 40 pg treatments of patient-derived tau, seeding rates were much more stable (Figure 4.E, F). In mixed effects models, the *vcov*_rep_ converged on 0, indicating the variance due to replicate was now technically negligible (Supplementary Table 5). We observed the same pattern of seeding rates across technical replicates at 40 pg, with no significant differences between technical replicates (Figure 4.F). However, one PiD case consistently resulted in higher seeding than others, despite normalised amounts of tau being used, indicating potential patient-specific differences in seeding capacity. Similar to 15 pg tau treatments, the aggregate size was stable across replicates (Figure 4.G), suggesting most of the variation is limited to seeding rate. In contrast to the 15 pg treatment, 40 pg additions of tau resulted in more consistent transfection efficiencies over 3 separate replicates (Figure 4.D, H).

### Patient-derived tau has disease-specific impacts on autophagy-lysosomal pathways

We next examined how patient-derived tau seeds affect the autophagy-lysosomal system. We conducted immunofluorescence of established markers of lysosomal proteolytic activity (CTSD), lysosomal morphology and abundance (LAMP1), lysosomal membrane damage (LGALS3) and autophagic flux disruption (SQSTM1/p62) on cells treated with 40 pg/well of tau (Figure 5.A). We observed a strong reduction in CTSD-positive puncta per cell in cells seeded with AD- and PiD-derived tau compared to PSP-derived tau (adjusted p-value: PiD-AD = 0.014, PSP-AD = 0.001, PSP-PiD = 1.07 × 10^-8^, Figure 5.A, B). This effect was most pronounced in PiD-treated cells, where CTSD was almost completely absent outside of regions co-localising with tau aggregates, indicating a profound sequestration of active lysosomal proteases to PiD-derived tau. In all cases treated with tau, we observed colocalization of CTSD with a subset of tau puncta (Figure 5.A). In contrast, cells treated with PSP-derived tau had increased levels of CTSD compared to other patient-derived tau treatments. The chloroquine-treated positive control cells had drastically reduced CTSD puncta (Supplementary Figure 4).

**Figure 5.**
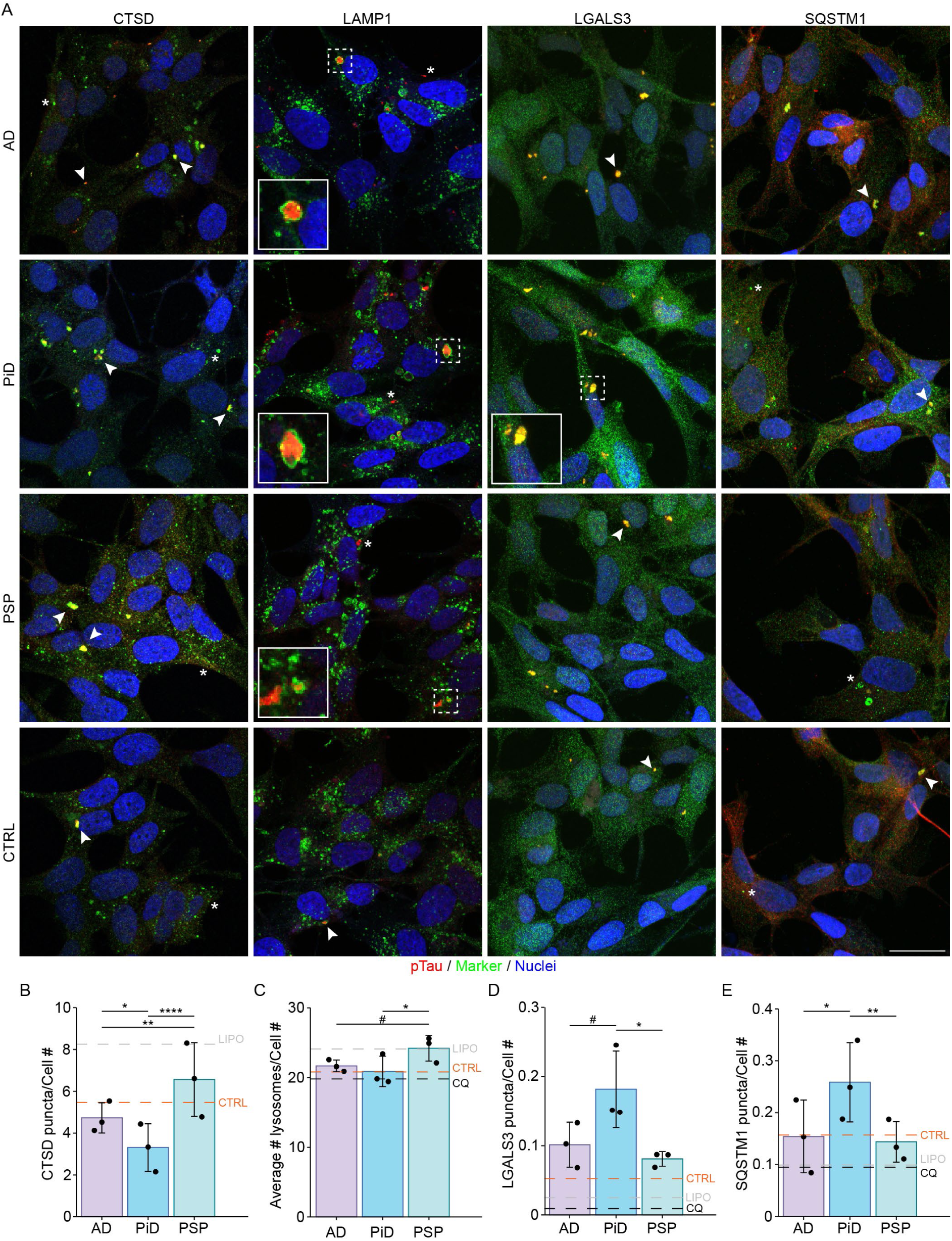
Patient-derived tau has disease-specific impacts on the autophagy-lysosomal pathway. Representative confocal images of SH-SY5Y cells treated with 40 pg of patient-derived tau from AD, PiD, PSP and controls. A) AT8-immunoreactive pTau (red) is co-stained with CTSD, LAMP1, LGALS3 or SQSTM1 (green). B) Quantification of the impact of patient-derived tau seeds on CTSD puncta per cell. CQ treatment omitted due to quantification artefacts. One-way ANOVA with Tukey’s post-hoc test. Adjusted p-values: PiD-AD = 0.014, PSP-AD = 0.001, PSP-PiD = 1.07×10^-8^. C) Quantification of tau seeds’ impact on the number of LAMP1-positive lysosomes per cell. One-way ANOVA with Tukey’s post-hoc test. Adjusted p-values: PiD-AD = 0.73, PSP-AD = 0.063, PSP-PiD = 0.009. D) Quantification of the impact of patient-derived tau seeds on LGALS3-positive puncta per cell. One-way ANOVA with Tukey’s post-hoc test. Adjusted p-values: PiD-AD = 0.088, PSP-AD = 0.79, PSP-PiD = 0.038. E) Quantification of the impact of patient-derived tau seeds on SQSTM1 puncta per cell. One-way ANOVA with Tukey’s post-hoc test. Adjusted p-values: PiD-AD = 0.011, PSP-AD = 0.95, PSP-PiD = 0.005. Scale bar = 20 μm. *N =* 3 biological replicates for AD, PiD and PSP, *n =* 1 CTRL. CQ = chloroquine treatment. Tau-free chloroquine and lipofectamine controls are shown in Supplementary Figure 4. Arrows indicate tau pathology colocalised with the marker. * Indicate no colocalisation with the marker.

The number of LAMP1-positive vesicles and the total LAMP1-positive area were significantly increased in PSP-derived tau treated cells (adjusted p-value count: PSP-AD = 0.06, PSP-PiD = 0.009; area: PSP-AD = 0.002, PSP-PiD = 0.009; Figure 5.A, C, Supplementary Figure 5), consistent with enhanced lysosomal biogenesis in response to PSP tau. In all tau-seeded conditions, LAMP1 signal frequently encased large aggregates or appeared as membrane-associated puncta containing invaginations (Figure 5.A insets). Notably, in all cases a substantial proportion of tau aggregates were not enclosed in LAMP1-positive vesicles, suggesting escape from the lysosomal system (Figure 5.A).

PiD-derived tau treated cells had significantly increased LGALS3-positive puncta compared to PSP (adjusted p-value: PSP-PiD = 0.04). This indicates robust induction of lysosomal membrane damage by PiD-derived tau (Figure 5.A, D). By 4 days post-treatment, LGALS3-positive puncta were frequently clustering near the cell surface or in blebs protruding from the cell (Figure 5.A inset), consistent with lysosomal exocytosis. Both AD- and PSP-derived tau treated cells had similar degrees of LGALS3 puncta formation. Whilst this was similar in count to control-derived tau treated cells, control tau treated LGALS3 puncta were often small in appearance. Neither chloroquine nor lipofectamine-3000 elicited any LGALS3 puncta formation (Supplementary Figure 4).

Finally, autophagic flux disruption was assessed by SQSTM1 puncta count. This was most prominent in PiD-derived tau treated cells. SQSTM1-positive puncta were significantly increased in response to PiD-derived tau compared to AD- or PSP-derived tau (adjusted p-value: PiD-AD = 0.01, PSP-PiD = 0.005; Figure 5.A, E), suggesting PiD-derived tau has a robust and direct impact on autophagy. SQSTM1 was co-localised with a subset of tau aggregates in both PiD- and AD-derived tau treated cells, but rarely in PSP-derived tau treated cells. Neither chloroquine nor lipofectamine-3000 elicited strong SQSTM1 puncta formation (Supplementary Figure 4).

Collectively, these findings reveal tauopathy-specific perturbations of the autophagy-lysosomal systems. AD- and PiD-derived tau rapidly decreased lysosomal protease content, whilst PiD-derived tau uniquely induced lysosomal damage and impaired autophagic flux. In contrast, PSP-derived tau had minimal impact on lysosomal function but was associated with increased lysosomal abundance and active proteases, implying a biogenetic response. These differences occurred despite seeding with the same concentration of tau from each disease and very similar presence of background proteins, suggesting these effects reflect intrinsic disease-specific properties of these tau seeds.

### Patient-derived tau can seed multiple cell lines

To determine the transferability of this protocol to other cell types, we tested tau seeding in three other immortalised cell lines. We used MO3.13 cells (a hybrid cell derived from fusing rhabdomyosarcoma cells with human oligodendrocytes), U-87s (glioblastoma cells with an epithelial phenotype) and U-118s (glioblastoma cells with a mixed phenotype [bipolar epithelial, stellate-glial and epithelioid]). These cell lines were treated with 40 pg of patient-derived tau and imaged 4 days post-transfection.

Overall, we observed similar seeding patterns to SH-SY5Y cells in all three lines for each disease (Figure 6). However, all three cell lines had reduced tau seeding rates compared to what was observed for SH-SY5Y cells, typically about half the seeding rate of SH-SY5Y cells (Figure 3.A, C and Figure 6.A-D). All cell lines were most robustly seeded by PiD-derived tau (Figure 6.B-D), similar to SH-SY5Y cells. However, in both MO3.13 and U-118 cell lines, we observed a stronger bias in tau aggregate distribution, often with epithelial-like cells having few aggregates, whilst more stellate cells (glial-like appearance) had the most aggregates. U-118s had a high degree of endogenous punctate tau expression, inflating some of the puncta counts acquired for this cell line (Figure 6.A, D). U-87s had markedly fewer tau aggregates detected per cell, possibly due to their higher proportion of epithelial-like cells. Overall, these results showed that tau aggregates can seed multiple cell lines in the same pattern (PiD-derived tau seeds being most robust at seeding), but cell type and phenotype likely impact seeding competency.

**Figure 6.**
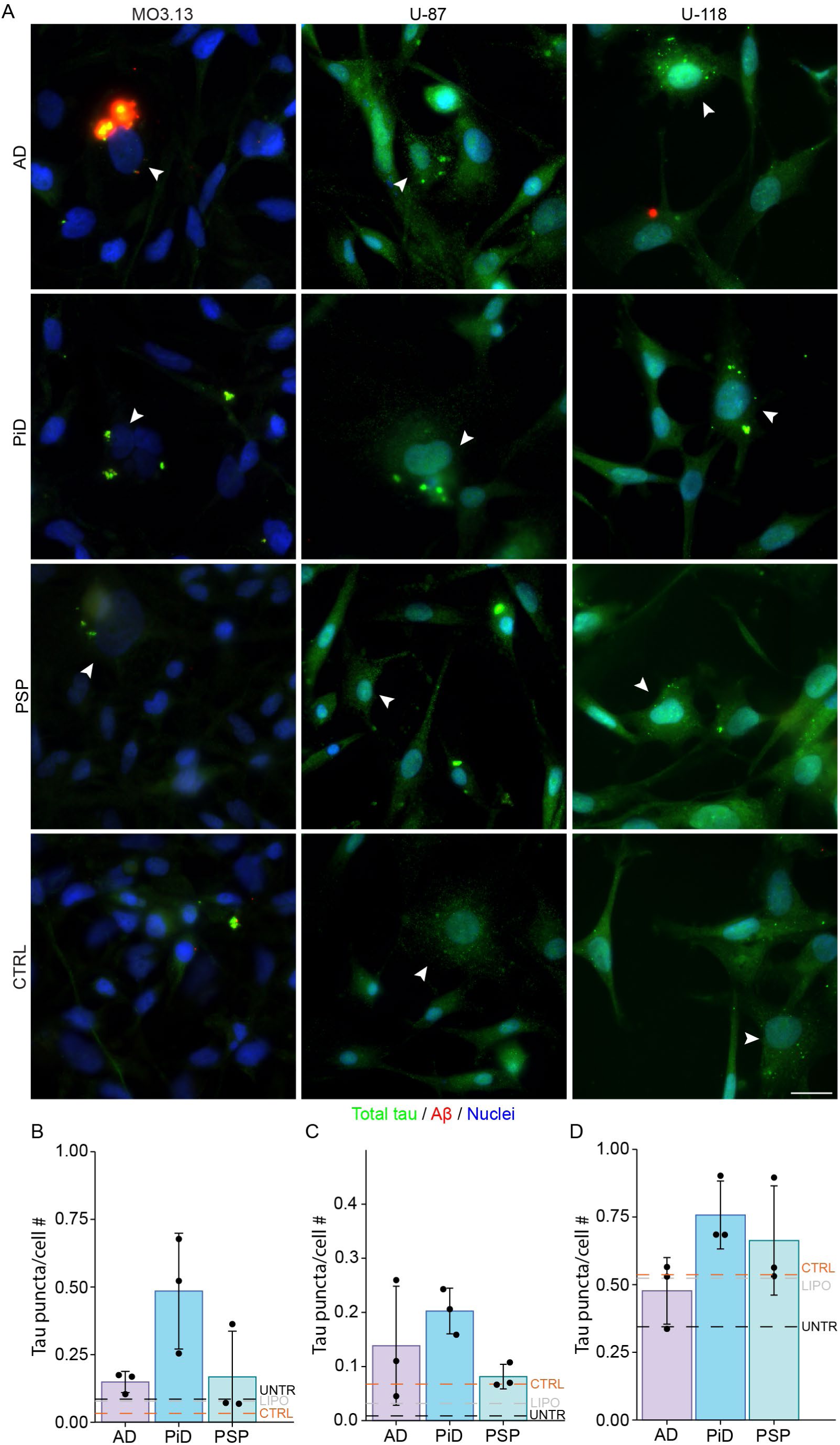
Tau and Aβ seeding patterns in MO3.13, U-87 and U-118 cells. A) Representative images of MO3.13, U-87 and U-118 cells treated with patient-derived tau seeds from AD, PiD, PSP and CTRL samples. B-D quantification of the number tau aggregates normalised to the number of cells induced by patient-derived tau seeds in B) MO3.13 cells, Kruskal-Wallis’ test p-value = 0.182. C) U-87 cells, One-way ANOVA group-wide p-value = 0.209. D) U-118 cells, One-way ANOVA group-wide p-value = 0.263. Scale bar = 20 μm.

### Patient-derived tau can seed iPSC-derived astrocytes

In mixed phenotype cell lines, we observed a preponderance of tau aggregates in stellate (glial-like) cells. To determine if patient-derived tau aggregates could readily seed more complex cell systems, and if a purer mix of glial cells would demonstrate the same uptake of tau aggregates, we used the same protocol to transfect iPSC-derived astrocytes (iAstrocytes). We seeded iAstrocytes with 40 pg of tau from each biological case and imaged these 4 days after treatment (Figure 7.A, B). We observed extremely robust seeding of tau aggregates per cell, with an average of 9.2 ± 1.5 for AD-derived tau and 6.6 ± 2.5 for PiD-derived tau (Figure 7.A, B, C). PSP- and control-derived tau seeded at a much lower rate, 1.8 ± 1.2 and 0.8 tau puncta/cell number, respectively (Figure 7.C). In all instances, this seeding outperformed the tau seeding seen in SH-SY5Y cells by approximately 25x in AD-, 7x in PiD- and 3.6x in PSP-derived tau treated cells (Figure 3.A, C, Figure 7.C). Overall, we observed a high seeding efficiency of iAstrocytes with AD and PiD patient-derived tau, with tau puncta observed in 50.6% of AD-treated cells and 50.8% of PiD-treated cells (Figure 7.D). In contrast, a lower seeding efficiency was observed for PSP patient-derived tau, with 31.6% of cells having tau puncta (Figure 7.D). These disease-specific seeding efficiencies were particularly notable given that cells were seeded with the same amount of tau.

**Figure 7.**
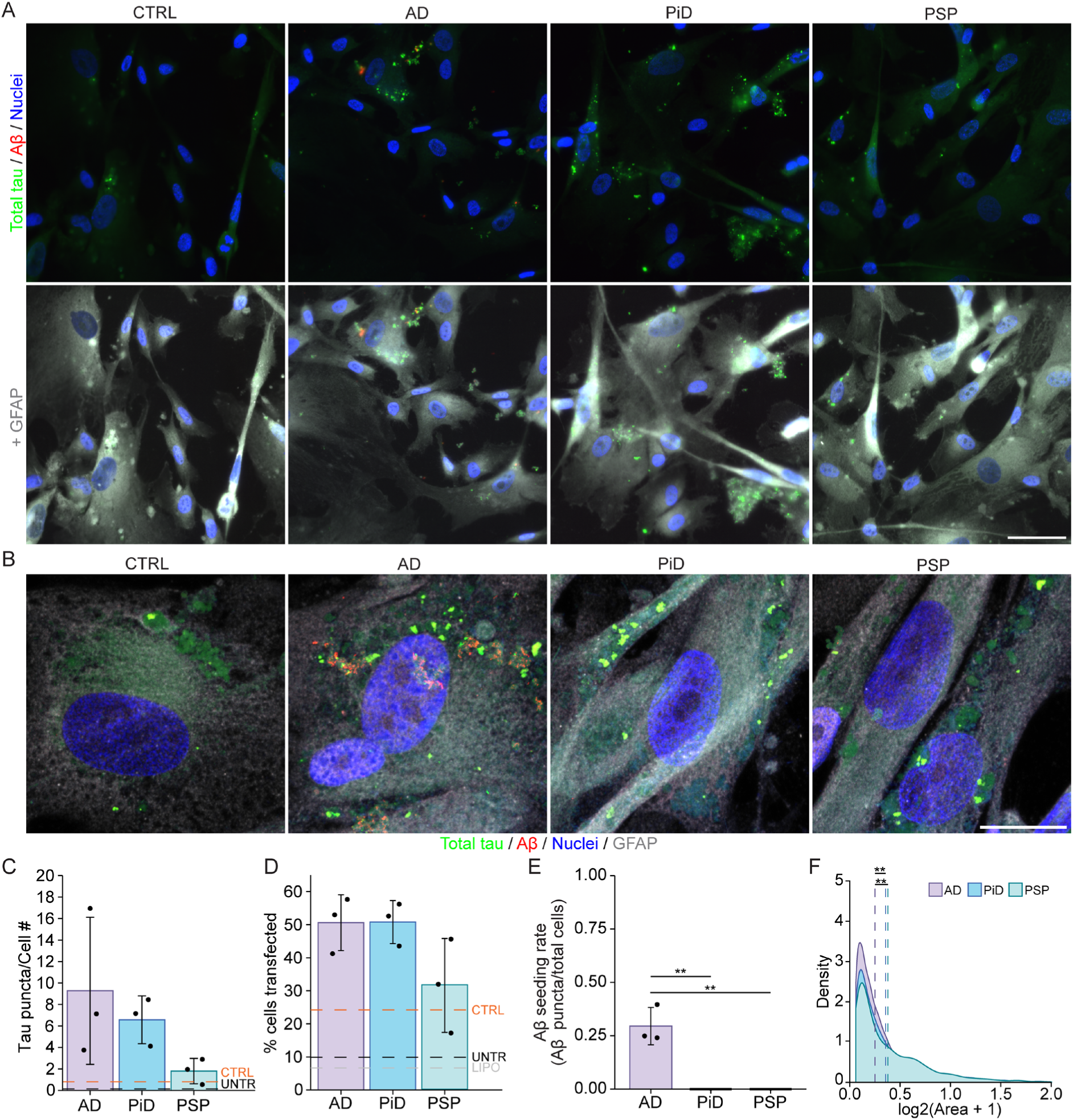
Patient-derived tau seeds robustly induce pathology in iAstrocytes. A) Representative widefield images of tau and Aβ pathology in iAstrocytes treated with patient-derived tau seeds. Scale bar = 50 μm. B) Representative cropped confocal images of tau and Aβ pathology in iAstrocytes. Images are maximum projections. Scale bar = 20 μm. C) Quantification of tau aggregate counts per cell. One-way ANOVA group-wide p-value = 0.144. D) Quantification of tau seeding efficiency. One-way ANOVA group-wide p-value = 0.1. E) Distribution of tau aggregate sizes in iAstrocytes between AD, PiD and PSP. One-way ANOVA with Tukey’s post-hoc test. Adjusted p-values: PiD-AD = 0.005, PSP-AD = 0.002, PSP-PiD = 0.60. F) Quantification of Aβ aggregate counts per cell. One-way ANOVA with Tukey’s post-hoc test. Adjusted p-values: PiD-AD = 0.001, PSP-AD = 0.001, PSP-PiD = 1.0.

Consistent with other seeding experiments, we only observed Aβ aggregates in cells treated with AD-derived tau seeds (Figure 7.E). Interestingly, in iAstrocytes, Aβ and tau puncta were usually located at the extreme edges of the cell in what appeared to be large blebbing bodies (Supplementary Figure 6). Furthermore, these aggregates were on average much smaller than those observed in SH-SY5Y cells (all SH-SY5Y tau puncta average size = ∼1.58 ± 0.59 µm^2^. iAstrocytes average size [µm^2^]: AD 0.25 ± 0.02, PiD 0.36 ± 0.04, PSP 0.38 ± 0.01) (Figure 3.E, Figure 7.F).

## Discussion

Here, we present a robust and reproducible patient-derived tau seeding model without the need for tau overexpression. Critically, our model reproduces the complex tau pathology observed in sporadic AD and sporadic primary tauopathies, including phosphorylation and isoform biases. Sporadic cell models are needed given that most clinical cases are sporadic, but most current models rely on mutated or synthetic tau that is different to that observed in sporadic disease. Our results demonstrate that tau from different tauopathies has significantly different seeding propensity and downstream effects on cellular pathways. These differences exist despite seeding the same amount of tau. This highlights the unique effect different strains of tau have in disease. Furthermore, we demonstrate that our model is transferable to multiple cell lines, including several immortalised cell lines and iPSC-derived cells, providing flexibility for future studies. Another key aspect of our models is that our AD-derived seeds have a mix of both Aβ and tau pathology, allowing for the unique opportunity to study the synergistic effects of the two pathologies on cellular processes and more accurately reflect human disease [38, 39].

Our data show disease-specific differences in tau-seeding rates that have been previously observed in bio-sensor models [51, 52]. Our data highlight that this occurs without overexpression of tau constructs, indicating that this is likely a genuine human disease-specific effect. PiD-treated cells typically displayed the greatest rate of seeding. This is partially supported by biosensor cell lines where PiD-seeds aggregate at the same or higher rates as AD in HEK-293 cells expressing 3R tau [26, 30]. Curiously, whilst AD-derived tau had the highest enrichment of tau (and co-enriched Aβ) in the mass-spectrometry results, it did not have a concomitantly high seeding rate. Furthermore, Aβ seeding was highly variable, perhaps reflecting broad differences in concentration between AD patients, a higher degree of inertness of Aβ aggregates or a result of normalizing seeding concentration to tau quantity.

PiD-derived tau resulted in the highest rate of seeding in all immortalised cell lines. However, contrary to expectations, in the iAstrocytes, AD and PiD were seeded at equivalent rates and transfection efficiencies. This coincided with much smaller aggregate sizes observed in iAstrocytes compared to SH-SY5Y cells. These results highlight cell-specific responses to tau seeds that are important to account for and explore in future studies. In contrast, PSP-derived seeds generally resulted in lower seeding across all cell types. This was particularly surprising in the iAstrocytes, given that tufted astrocyte tau pathology is a hallmark of PSP [19, 24, 38, 39]. However, the seeding rate for PSP-derived tau in iAstrocytes was approximately double that of SH-SY5Y cells. This may reflect less pathological activity from PSP-derived 4R-tau seeds, as observed in mouse models comparing AD and PSP [19, 24, 38, 39].

Autophagy-lysosomal disruption and damage are well-known features of neurodegeneration and can be directly triggered by Aβ and tau aggregates [17, 21, 53–57]. However, the exact mechanism of disruption triggered by specific tau strains is not yet understood. Here, as a proof-of-principle study, we investigated common autophagy-lysosomal markers to determine whether this model can detect disease-specific effects. PiD-derived tau had the most marked effect on autophagy and lysosomal activity, leading to increased LGALS3 and SQSTM1 puncta and depleted CTSD. Together, these indicate rapid and acute disruption of autophagosome formation, induction of lysosomal damage and reduction in lysosomal proteases. These changes stand in stark contrast to PSP-derived tau, which had minimal effects on autophagy or lysosomal damage but showed increased CTSD puncta and lysosomal number compared to PiD- and AD-derived tau. These changes in PSP-treated cells may indicate increased lysosomal biogenesis or limited impacts of PSP-derived tau on the lysosomal system. AD-treated cells tended to sit in the middle ground. AD-treated cells displayed similar levels of lysosomal damage as PiD-treated cells. But they only showed a significant difference in CTSD puncta, sitting between PiD- and PSP-treated cells. This was unexpected as tau and Aβ are thought to act synergistically to impair the lysosome [56–58]. This may indicate a few possible explanations: 1) PiD-derived tau is more disruptive of the lysosome, 2) Aβ derived by PTA precipitation is less pathologically active than in the brain, 3) Aβ affects other specific lysosomal processes not assessed here.

There are several limitations of this model. This includes the nature of human-derived protein preparations, including multiple contaminating proteins including several that were enriched at a higher concentration than tau or Aβ. We have controlled for this in the current study by inclusion of control patient-derived preparations in all experiments, which we show closely mimic the enrichment of background proteins. Seeding experiments must control for these background proteins to allow a genuine appraisal of tau-specific effects. While the presence of these additional proteins may be viewed as a technical limitation, we propose that accurately reflects the complexity of human AD and primary tauopathies in our model. Secondly, we used ELISA to measure the concentration of tau seeds, which may only readily assess soluble protein, leading to underestimates of total tau if the PTA pellets are not fully solubilised. Our results suggest that seeding experiments can be inherently variable, and we demonstrate here that there is a floor at which seeding becomes overly noisy and loses reproducibility. Furthermore, we observe cell-line-specific sensitivity to tau seeding. Future experiments need to assess that floor for their individual cell types, as this was only performed for SH-SY5Y cells in the current study. Indeed, future studies could apply this seeding model in full to iNeurons, complex mixtures of iPSC-derived cells or organoids where limited success has been achieved so far [50, 59]. Here, we have used Lipofectamine-3000 to facilitate seeding, which may alter tau (and Aβ in AD-derived preparations) uptake pathways in cells compared to physiological pathways. The removal of lipofectamine in future would be worth pursuing to further reduce non-physiological inputs in this model. The PTA protocol may remove some soluble but seed-competent species of tau, leading to an incomplete representation of tau species from each disease.

In summary, this study establishes a robust and physiologically relevant model of tauopathies, using patient-derived tau seeds with preserved biochemical properties from AD, PiD, PSP and control cases. By avoiding tau overexpression and controlling for contaminating proteins, this model allows direct comparison of tau protein uptake and downstream pathway disruptions driven by AD-, PiD- and PSP-derived tau. By reducing extrinsic manipulations, we leave room for further manipulation of the cell (e.g. overexpression or knock-down of interacting proteins). Our findings demonstrate disease-specific, acute impacts on the autophagy-lysosomal pathways, with PiD-derived tau exhibiting broad disruptions of the system, AD-derived tau disrupting active lysosomal peptidases and PSP-derived tau leading to compensatory lysosomal biogenesis. The transferability of this model across multiple neural and glial cell types, including human iAstrocytes, provides a powerful foundation for mechanistic studies of tau-cell interactions, identification of disease-specific modifiers, and medium-throughput therapeutic screening. This model offers a flexible and scalable framework for dissecting the cellular mechanisms of diverse tau strains and advancing the development of targeted therapies for tauopathies.

---

**Supplementary Figure 1** Dotblots of total homogenate, S1, P fractions derived by the PTA enrichment and control protein (BSA or milk protein) probed for total tau, pTau217, pTau396 [PHF13], pTau422 and oligomeric tau [T-22].

**Supplementary Figure 2** Volcano plots representing differentially enriched proteins between AD – PiD, AD – PSP and PiD – PSP contrasts. Differentially enriched proteins were determined by an FDR < 0.05 (horizontal dotted line) and |log_2_(fold change) | > 0.58 (vertical dotted lines).

**Supplementary Figure 3** A) Quantification of Aβ seed number dose-response of tau preparations derived from each condition. Line plot represents mean ± SEM. B) Representative confocal images of untreated (UNTX) and lipofectamine only (LIPO) cells.

**Supplementary Figure 4** Representative confocal images of lipofectamine and chloroquine treated SH-SY5Y cells. Tau (red) is co-stained with CTSD, LAMP1, LGALS3 or SQSTM1 (green).

**Supplementary Figure 5** Plot of normalised LAMP1-positive area normalised to cell count per field. One-way ANOVA with Tukey’s post-hoc test. Adjusted p-values: PiD-AD = 0.95, PSP-AD = 0.0022, PSP-PiD = 0.00088.

**Supplementary Figure 6** Confocal z-stack of iAstrocytes treated with 40 pg of AD-derived tau. Image taken in 1024×1024 pixels at 204 nm pixel size. Z-stacks taken at 0.125 µm. Blue labels nuclei, green labels tau, red Aβ and grey labels the cell membranes (CellMask Far Red).

## Data Availability

The mass spectrometric raw files are accessible at https://massive.ucsd.edu under accession MassIVE MSV0000XXXXX. Processed mass spectrometry results are included in the supplementary files of this paper. All other data is available from the corresponding author on reasonable request.

## Acknowledgments

The authors acknowledge the technical and scientific assistance of Sydney Microscopy & Microanalysis, the University of Sydney node of Microscopy Australia. Human brain tissues were received from the Sydney Brain Bank which is supported by Neuroscience Research Australia. We acknowledge the Cedars-Sinai Medical Centre’s David and Janet Polak Foundation Stem Cell Core Laboratory for providing the iPSC line used within the study.

## Declarations

This research project was approved by the Human Research Ethics Committee of the University of Sydney and complies with the statement on human experimentation issued by the National Health and Medical Research Council of Australia. Tissues were selected from a neuropathological series collected by the Sydney Brain Bank through regional brain donor programs in Sydney, Australia. The brain donor programs hold approval from the Human Research Ethics Committees of the South Eastern Sydney Area Health Services and comply with the statement on human experimentation issued by the National Health and Medical Research Council of Australia.

## Authors’ contributions

ED and TK conceived the experiments. TK and AS, KB, SG, and CA performed the experiments. EK and BU ran and processed the LC-MS and searches. TK, AS, and JG performed the analysis and made the figures. G.H. provided neuropathological assessment of cases and expert advice about the interpretation of the data. ED, GH, EW and MK obtained funding for the project. TK wrote the paper. All authors approved the final manuscript.

