## Supplementary Figures 1-5 for "Seeding patient-derived tau induces tauopathy-specific aggregation and lysosomal disruption in human cells"

Supplementary Figure 1

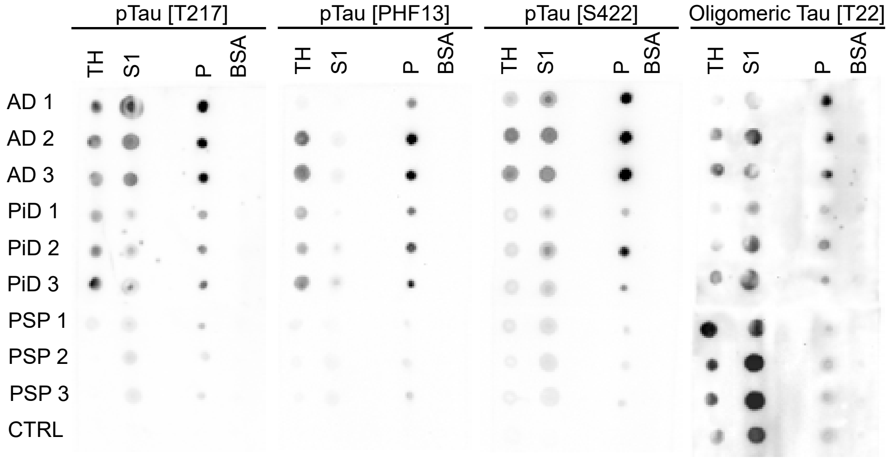

Supplementary Figure 2

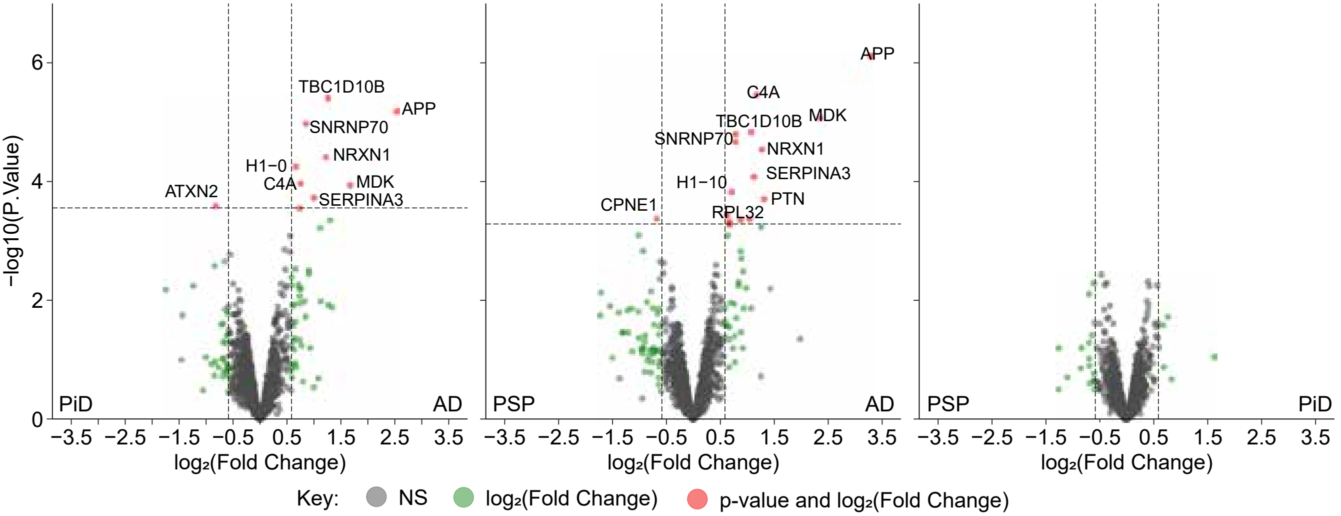

Supplementary Figure 3

A

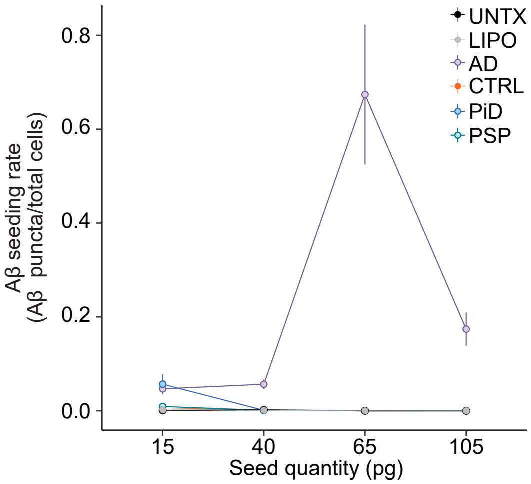


B

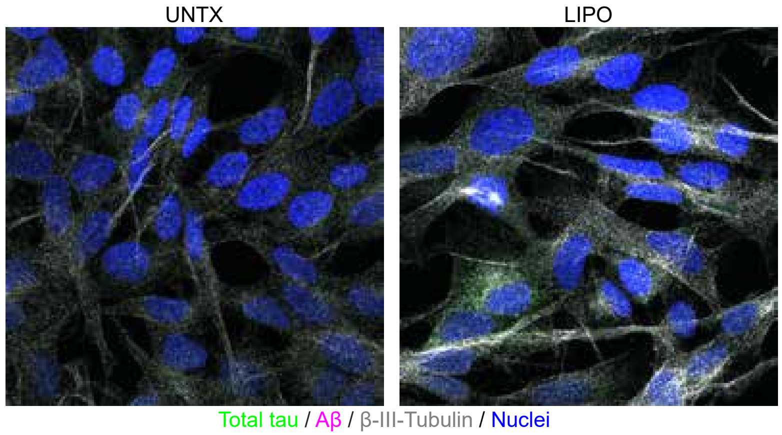

Supplementary Figure 4

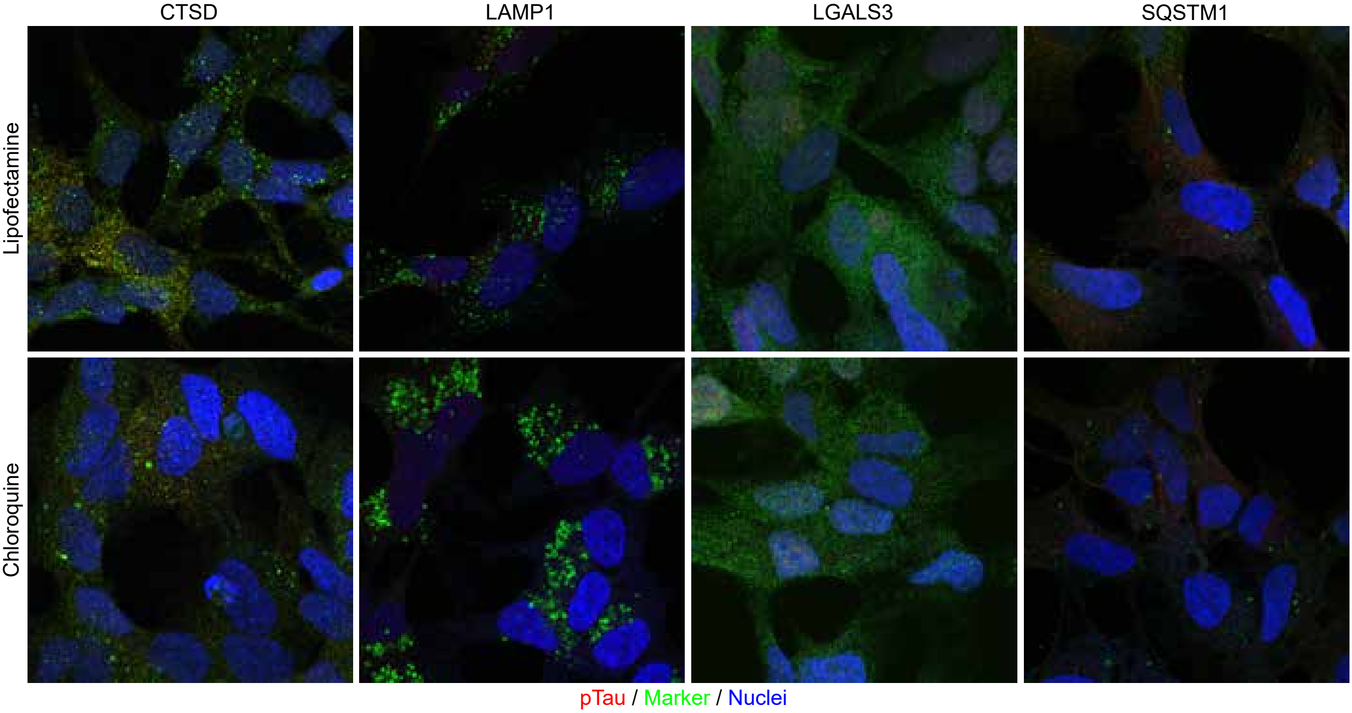

Supplementary Figure 5

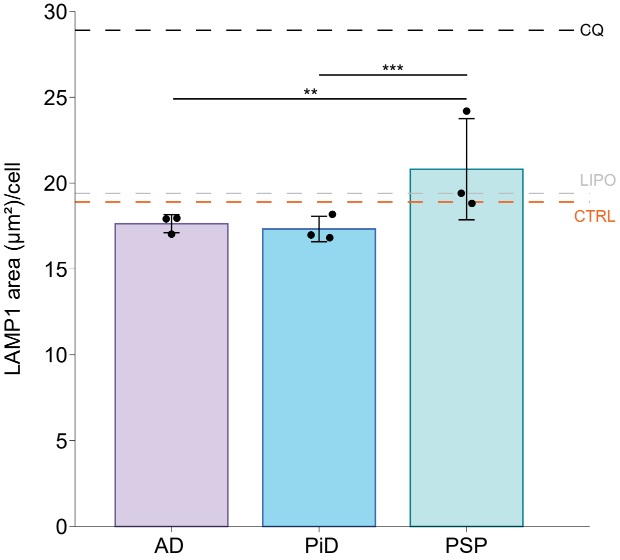
