## Supplementary figures and images for "Seeding patient-derived tau induces tauopathy-specific aggregation and lysosomal disruption in human cells"

### Supplementary Figure 6

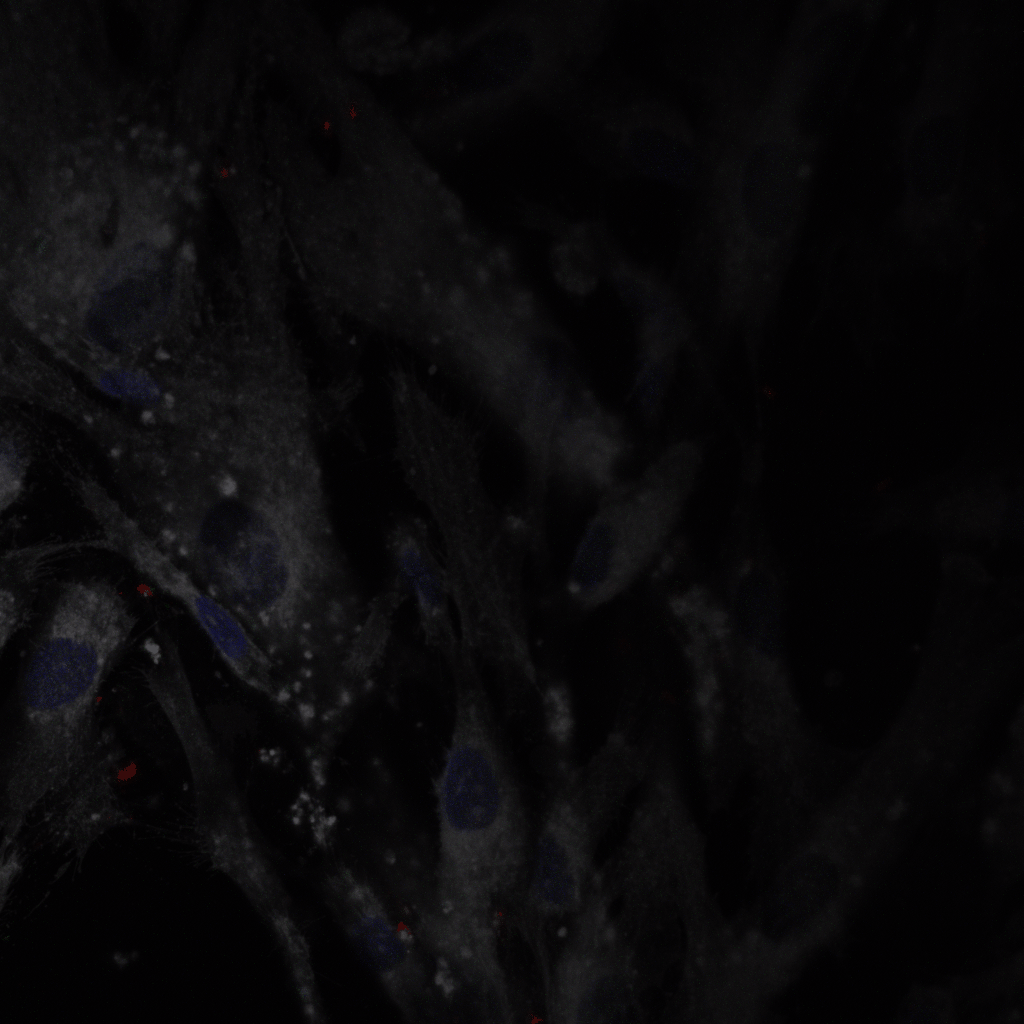
